# Cultivation and Fluorescent in situ hybridization suggest that some shipworm species acquire endosymbiotic bacteria through indirect horizontal transmission

**DOI:** 10.1101/2022.11.13.516348

**Authors:** Lauren Speare, Daniel L Distel

## Abstract

Beneficial microbial symbionts provide essential functions for their host from nutrients to defense against disease. Whether hosts acquire their symbionts directly from parents (vertical transmission) or by sampling from the environment (horizontal transmission) can have dramatic impacts on host adaptability and, in the case of ecosystem engineers, ecosystem health. Wood-boring bivalve mollusks (Teredinidae shipworms) act as ecosystem engineers in marine environments, creating habitat out of submerged wood for fish and invertebrates. Essential to shipworm success is their community of endosymbiotic gill bacteria that produce the enzymes necessary for wood digestion. How shipworms acquire their symbionts, however, remains largely unexplored. Using culturing, fluorescence *in-situ* hybridization, confocal microscopy, and tank experiments, we provide evidence suggesting the mode of symbiont transmission the shipworms for either the shipworm, *Lyrodus pedicellatus* or *Teredo bartschi* or both. Symbiotic bacteria were not detected by cultivation or microscopy in brooding larvae within gravid adults or as veliger larvae collected from the water column, but were observed in adult specimens and juveniles that had begun burrowing into wood. These data suggest that the specimens examined have both aposymbiotic and symbiotic life phases and acquire their symbionts through indirect horizontal transmission. Our findings reveal how the long-term brooders *L. pedicellatus* and/or *T. bartschi* acquire their gill endosymbionts.

**IMPORTANCE:** How eukaryotic hosts acquire their microbial symbionts can have significant consequences for their ability to adapt to varied environments. Although wood-boring bivalve shipworms have diverse reproductive strategies and are found in unique environments across the globe, little is known about how they transmit their essential gill endosymbionts. We used the closely related shipworms, *Lyrodus pedicellatus* and/or *Teredo bartschi* to study how these long-term brooding shipworms acquire their gill endosymbionts. Our work, unlike previous claims for the broadcast spawning species *Bankia setacae* which reportedly transmits its symbionts directly from parent to offspring, suggests that juvenile *L. pedicellatus* and/or *T. bartschi* acquire their symbionts through horizontal transmission rather than directly from their parents. This work reveals the mechanism by which some brooding shipworm species acquire their symbionts, adding to our limited understanding of intracellular symbiont transmission of Teredinidae.

## INTRODUCTION

Nutritional symbioses allow eukaryotic hosts to exploit otherwise unavailable food sources, expanding the niches available to both symbiotic partners [1]. Many marine ecosystem engineers such as scleractinian corals, hydrothermal vent tubeworms, some deep-sea mussels, and wood-boring shipworms rely on their symbiotic microbes to create structural habitats in their respective ecosystems [1–4]. How these mutualistic associations are established and able to persist across multiple generations has important implications for host fitness and ecosystem health.

Theoretical studies indicate that mutualisms strongly favor vertical transmission of symbionts, i.e., direct transmission of symbionts from parents prior to offspring release [5–7]. Vertical transmission guarantees the next generation has symbionts and their associated benefits and has been linked with more cooperative symbionts and reduced risk of host exploitation [6]. Horizontally transmitted symbioses in which offspring acquire symbionts from the environment in each subsequent generation, however, are wide spread and allow hosts to adapt to new conditions [5]. Hosts that acquire symbionts from the environment can select for partners that confer local advantage [8, 9] and avoid genetic bottlenecks due to reproductive isolation of symbionts through many host generations [6, 7]. In marine environments, transmission of microbial symbionts occurs predominantly horizontally, even in the case of intracellular symbionts [10, 11], likely due to the ability of symbionts to more easily travel from one host to another, as compared to terrestrial environments where desiccation and osmolarity issues are apparent [10, 12]. While the mode of symbiont transmission has been of keen interest for many ecosystem engineers [1, 13], little is known about how shipworms acquire their intracellular endosymbiotic bacteria.

Dubbed termites of the sea [14], shipworms are marine wood-boring bivalve mollusks from the family Teredinidae that use wood as a source of shelter and food [3, 15, 16]. Shipworms are among the most divergent and diverse bivalve families [17] however, how these ecologically important bivalves digest wood remains unknown. Shipworms harbor dense communities of intracellular gill symbionts that produce cellulolytic enzymes; similarly to the majority of wood-eating animals, these enzymes may allow for wood digestion by the host [2–4]. Cellulase-producing symbionts are localized within the gland of Deshayes, a specialized region within the host gill, rather than in the digestive tract, allowing the host to directly consume wood-derived sugars without competing with their symbionts [18, 19]. Recent work has revealed that many shipworm hosts harbor up to three species that make up the majority of the community, as well as a diverse group of low abundance taxa, although the identity and relative abundance of these taxa vary by host species [20].

To date, few studies have explored symbiont transmission in shipworms. Sipe et al [21] studied symbiont distribution in the shipworm *Bankia setacea* and detected symbiont 16S rRNA in the gill, ovary, and egg using PCR amplification, suggesting *B. setacae* acquires their symbionts through vertical transmission. However, whether the same is true for other shipworm species remains unknown. In this work, we explored the mode of symbiont transmission for representative long-term brooding shipworms. Unlike most intracellular symbionts, many shipworm gill symbionts share characteristics of free-living bacteria, do not appear to be obligate, and be cultivated *in vitro* [15, 22, 23], thus making the shipworm symbiosis an intriguing model to explore symbiont transmission. Further, a community of shipworms is currently maintained in aquaria at the Ocean Genome Legacy Center (OGL), allowing for experimental manipulation. Using a combination of Fluorescence *In-Situ* Hybridization and confocal microscopy, culturing, and tank experiments we aimed to describe the mode of symbiont transmission for shipworms in the colony maintained at OGL.

At the time that this experimental work was performed, it was thought that the OGL shipworm community was composed only of *Lyrodus pedicellatus*, however the community also appears to contain a second species, *Teredo bartschi*. For this reason, we cannot rule out the possibility that specimens, especially larvae and juveniles, of one or both species were included in the described experiments. However, because *L. pedicellatus* and *T. bartschi* are closely related, it is unlikely that this will alter the conclusions of this work. Nonetheless, we are currently working to verify the results described here for each of these shipworm species independently prior to peer review publication. Until that time, we present our results here, referring to the experimental specimens as members of this mixed species OGL colony.

## RESULTS

### Bacterial symbionts infect adult *L. pedicellatus* and/or *T. bartschi* gill tissue

To determine the mode of symbiont transmission in the shipworms in the OGL colony, we first sought to localize bacterial symbionts within adult gill tissue (Fig 1A). Fluorescence in situ hybridization (FISH) was performed on paraffin-embedded 5 um thick sections from whole adult specimens. Experiments were performed with a general bacterial probe (EUB338) that is complementary to a portion of 16S rRNA found in all bacteria, and control probe (NON338) which accounts for non-specific binding of EUB338 [24]. These probes were used previously to localize bacterial symbionts in shipworm tissue [19, 25, 26], including *L. pedicellatus* [18]. Specific fluorescent signals were observed in the gills of all adult specimens examined (Fig 1B and 1C, Fig S1). Signal from the gill appears to come primarily from the interlamellar junction (center of the body), rather than the gill filament (Fig 1B and 1C), which is consistent with the previously described location of endosymbiont containing bacteriocytes [18]. All of the adult specimens examined had sections of the gill that contained bright EUB338 signal, and sections without any visible signal. This phenomenon is most likely due to the spatial orientation of the specimen within the paraffin block prior to embedding, as shipworms tend to curl during fixation. Alternatively, variation in signal could indicate variability in the concentration of symbionts along the gill. Hybridization with the NON338 did not generate the florescent signals (Fig S1).

**Figure 1.**
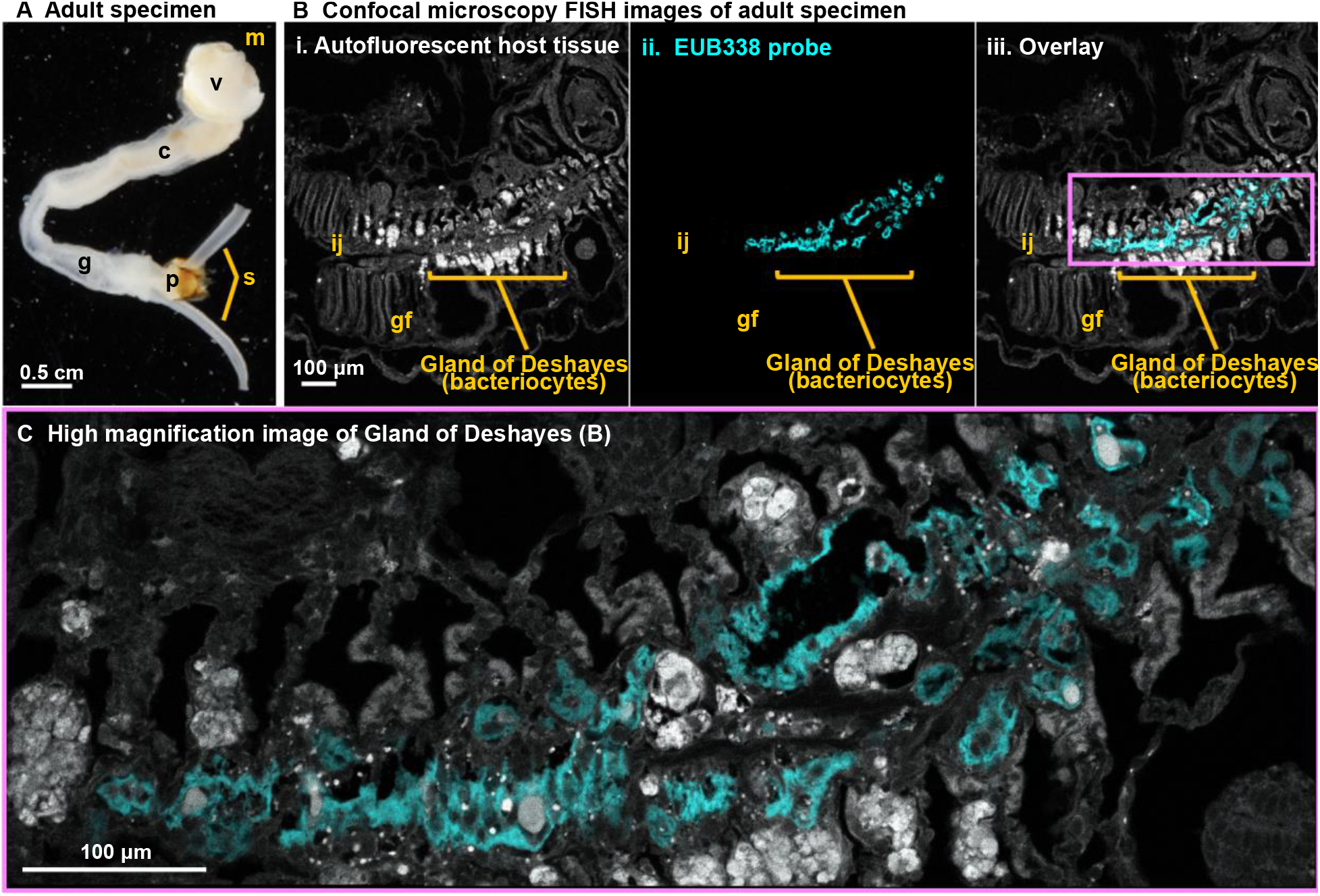
EUB338 can be used to detect intracellular bacteria within adult *L. pedicellatus* and/or *T. bartschi* Gland of Deshayes. (A) Adult shipworm specimen from the OGL colony. (B) Confocal microscopy FISH images of a representative adult specimen focused on the gill; each panel shows: i. autofluorescent host tissue, ii. signal from the eubacterial EUB338 probe, and iii. an overlay of i and ii. Control images with the non-specific NON338 probe are shown in supplemental information. (C) High magnification image of gland of Deshayes indicated by the pink box in Biii. Abbreviations are as follows: cecum (c), gill (g), gill filament (gf), interlamellar junction (ij), mouth (m), pallet (p), siphon (s), valve or shell (v).

These observations are consistent with previous studies and suggest that adult *L. pedicellatus* shipworms harbor symbiotic bacteria within their gill tissue [18]. We took two approaches to examine bacterial symbionts across *L. pedicellatus and/or T. bartschi* ontogeny, and thus determine the mode of symbiont transmission for the OGL shipworm colony: 1) quantify bacterial abundance through culturing, and 2) localize bacteria within shipworm tissue using FISH and confocal microscopy.

### Endosymbiotic bacteria can be cultured from juvenile yet not larval *L. pedicellatus and/or T. bartschi* shipworms

Although many endosymbiotic microbes remain unculturable, a portion of the *L. pedicellatus and/or T. bartschi* gill microbiome can be readily cultured adult specimens and importantly, *Teredinibacter turnerae*, is commonly isolated from adult *T. bartschi* specimens [15, 22, 23]. Therefore, to begin determining bacterial abundance in shipworms of different life stages, we quantified bacterial colony forming units (CFUs) from shipworm tissue using shipworm basal medium (SBM) that is selective for marine cellulolytic bacteria, including shipworm symbionts. Shipworms were collected from the OGL colony at the following life stages: burrowing juveniles, and developed juveniles were extracted from wood, veliger larvae were collected from the surface of wood or the water column, and brooding larvae were dissected from the gill tissue of gravid adults. Specimens were either rinsed with filter sterilized autoclaved sea water (FSASW) five times and dissected/homogenized immediately or flash frozen for 24 hours prior to dissection/homogenization. Brooding larvae and veliger larvae were pooled into groups of 3 larvae prior to homogenization. Tissue homogenate was then serially diluted and plated onto SBM agar plates to quantify bacterial abundance across ontogeny. CFUs were not detected in any brooding larvae regardless of sample preparation (Fig 2) and CFUs were detected at the limit of detection (1 CFU at 10⁰ dilution) in 2 out of 8 pools (6 of 24 larvae) of FSASW rinsed veliger larvae, yet not any frozen veliger larvae (Fig 2). Burrowing juveniles that had begun initial metamorphosis and minor burrowing into the wood contained approximately 10^5^ CFUs per animal, while more developed juveniles that had begun to elongate contained approximately 10^6^ CFUs per animal (Fig 2). There was no significant difference in the number of CFUs collected between only rinsed or rinsed and then frozen specimens at the life stages examined.

**Figure 2.**
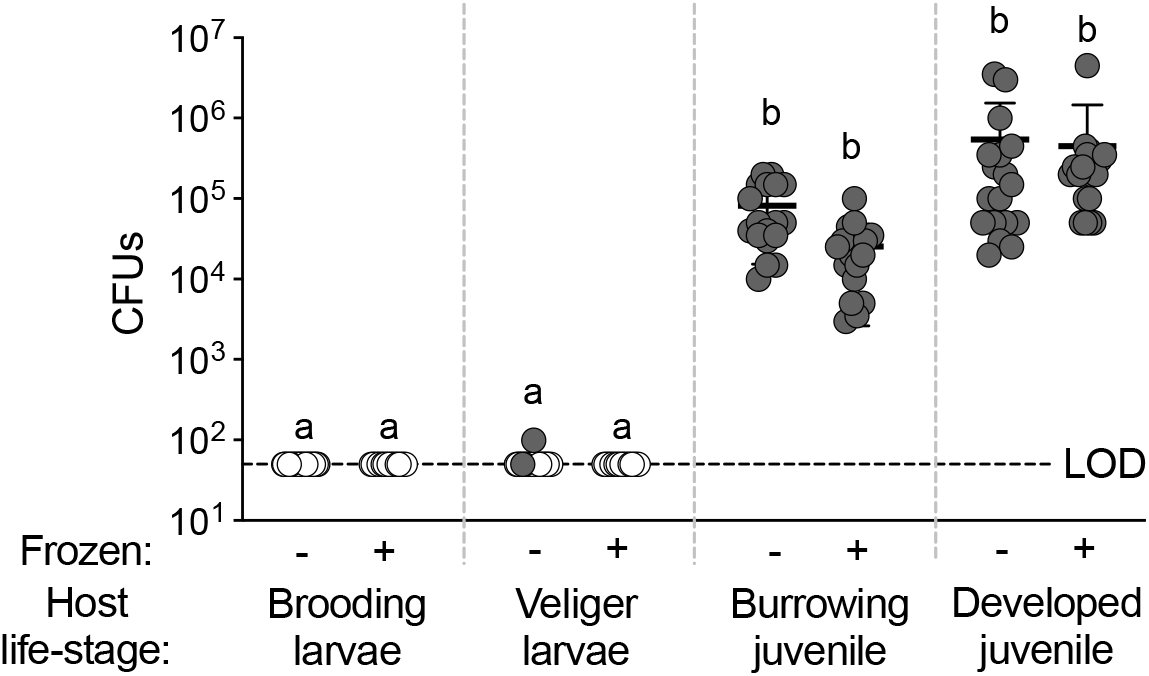
Bacteria can be cultivated from juvenile shipworms. Total CFUs from larvae that were rinsed with filter-sterilized and autoclaved sea water (−) or rinsed and then frozen for 24 hours (+) and plated onto SBM media. Host developmental stage is indicated on x-axis. Each circle represents CFUs for 1 (juvenile) or 3 (larvae) organisms. White circles indicate limit of detection (< 1 CFU from undiluted spot). Each experiment was performed twice with at least 8 biological replicates.

These data yield two important observations: 1) neither brooding nor veliger larvae contain a detectable level of endosymbiotic, culturable bacterial symbionts, and 2) juveniles that have begun go undergo metamorphosis (burrowing and more developed juveniles) contain a substantial community of culturable bacteria. These data are consistent with a hypothesis whereby *L. pedicellatus* and/or *T. bartschi* acquire endosymbiotic bacteria from the environment rather than directly from a parent during gestation.

### Symbionts can be visualized in juvenile *L. pedicellatus* and/or *T. bartschi* through FISH microscopy

We sought to test the hypothesis that *L. pedicellatus* and/or *T. bartschi* are aposymbiotic in early development by determining the spatial localization of bacterial symbionts within shipworm tissue using FISH and confocal microscopy. Given that our CFU data suggest that shipworms within the OGL colony do not acquire bacterial symbionts until they begin to burrow into wood, we predicted that we would not detect a bacterial signal in brooding or veliger larvae, and only observe a signal in more developed juveniles that had already begun to burrow. We first examined brooding larvae by imaging gravid adults. Although a signal was observed in adult gill lamellae, which is where brooding larvae reside, we were unable to detect a EUB338 signal from any brooding larvae themselves (Fig 3A, Fig S2) (n=34), suggesting that these larvae do not harbor endosymbiotic bacteria.

**Figure 3.**
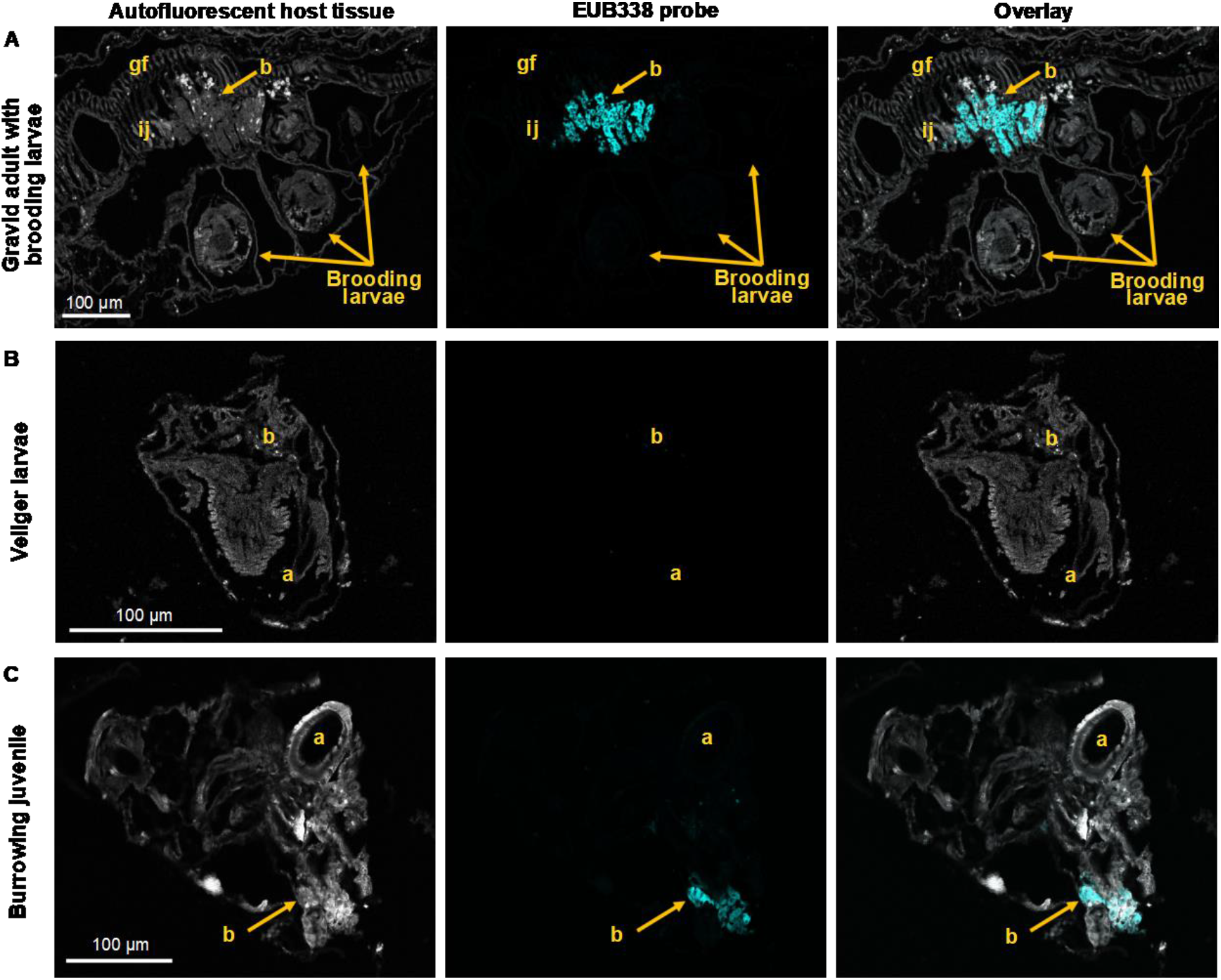
Bacteria are detected in burrowing juvenile shipworms via FISH. Confocal microscopy images of representative shipworms across ontogeny: A) brooding larvae within gravid adults, B) veliger larvae, and C) burrowing juveniles that have begun metamorphosis; each panel shows: autofluorescent host tissue (left, gray), signal from the eubacterial EUB338 probe (center, cyan), and an overlay of the two (right). Control images with the non-specific NON338 probe are shown in supplemental information. At least eight animals were imaged at each life stage. Abbreviations are as follows: adductor (a), bacteriocytes (b), gill filament (gf), interlamellar junction (ij), and pallet (p). Scale bars indicate 100 μm.

We next interrogated veliger larvae that were collected from the water column and the surface of wooden baits, and juvenile shipworms that had just begun to burrow into wooden baits for the presence of a EUB338 bacterial signal. Similar to the brooding larvae, no detectable signal was observed for veliger larvae (Fig 3B, Fig S3) (n=8). However, the majority of more developed larvae (8 out of 10) did contain a bright signal with the EUB388 probe in tissue resembling bacteriocytes within adult gill tissue (Fig 3C, Fig S4). Hybridization with the NON338 negative control probe did not generate a florescent signal (Fig S4). These data suggest that *L. pedicellatus* and/or *T. bartschi* have an aposymbiotic life stage, as brooding and veliger larvae, and acquire their symbionts by the time they have begun metamorphosis and burrowing.

### *L. pedicellatus* and/or *T. bartschi* larvae acquire bacteria through indirect horizontal transmission

Finally, we sought to determine the environmental source of horizontally acquired bacterial symbionts for juvenile *L. pedicellatus* and/or *T. bartschi* shipworms. Based on the CFU and FISH microscopy data, we predicted that juvenile shipworms acquire their symbionts either from the water column, or the wood on which the settle and burrow. We performed a series of tank experiments in which veliger larvae were collected from adult tanks, rinsed with FSASW, and allowed to settle on either sterile (autoclaved) or nonsterile wooden baits in either FSASW or larva-free water collected from tanks containing adult, larvae-producing *L. pedicellatus* and/or *T. bartschi*. We first tested whether larvae could settle and metamorphose in a “sterile” environment, on autoclaved wooden baits in FSASW, and did not observe any settlement or metamorphosis (n=93) (Table 1). We repeated this experiment with filtered water collected from adult tanks and similarly did not observe any settlement or metamorphosis on sterilized baits (n=94) or non-sterilized baits (n=76) (Table 1). We then repeated the experiment with adult tank water, non-sterilized baits, and an additional bait containing non-larvae producing adult shipworms. In this treatment, we observed settlement, burrowing, and initial metamorphosis for 13% of larvae (+/− 3% standard deviation) (n=78) (Table 1).

**Table 1.** Results of settlement experiments in sterile, semi-sterile, and standard conditions. Veliger larvae were collected from the water column of adult-containing tanks, rinsed with FSASW, and transferred to tank conditions indicated in the first column. Three to four trials of each experiment were performed with at least 21 veliger larvae per trial (total larvae n=341).

| Tank Conditions | Trial # | # veliger larvae | # burrowing juveniles |
| --- | --- | --- | --- |
| Sterilized baits in FSASW | 1 | 25 | 0 |
|  | 2 | 22 | 0 |
|  | 3 | 21 | 0 |
|  | 4 | 25 | 0 |
| Sterilized baits in adult tank water | 1 | 25 | 0 |
|  | 2 | 23 | 0 |
|  | 3 | 22 | 0 |
|  | 4 | 24 | 0 |
| Non-sterilized baits in adult tank water | 1 | 25 | 0 |
|  | 2 | 25 | 0 |
|  | 3 | 26 | 0 |
| Non-sterilized baits in adult tank water with non-larvae producing adults | 1 | 25 | 4 |
|  | 2 | 26 | 3 |
|  | 3 | 27 | 3 |

**Table 2.** Number of *L. pedicellatus* specimens examined via FISH and confocal microscopy. The number of specimens with a detectable EUB338 signal are indicated in the right column; at least two separate experiments were performed for animals at each life stage.

| Life Stage | # of specimens | EUB338 signal |
| --- | --- | --- |
| Adult | 4 | 4 |
| Brooding larvae | 34 | 0 |
| Veliger larvae | 8 | 0 |
| Burrowing juvenile | 10 | 8 |

Results of these tank experiments yield several important findings: 1) *L. pedicellatus* and/or *T. bartschi* larvae are unable to settle and burrow into wood in a sterile environment, 2) larvae-free water from adult tanks does not induce burrowing, and 3) presence of adult shipworms, whether larvae-producing or not, promotes larval settlement. These findings combined with our culture and FISH microscopy data support a hypothesis whereby *L. pedicellatus* and/or *T. bartschi* have an aposymbiotic life stage during early development and horizontally acquire their symbionts prior to or during settlement and early metamorphosis.

## DISCUSSION

In this work, we sought to determine the mode of transmission for the widely-distributed shipworms, *L. pedicellatus* and/or *T. bartschi*. Previous work on this model shipworm revealed they harbor endosymbiotic, intracellular gill bacteria [15, 19]. These communities provide important cellulase and nitrogenase enzymes to their hosts [3, 15, 16, 27–29] and are relatively simple, composed of *Teredinibacter turnerae* and other gram-negative proteobacteria [15, 23]. Using a combination of culturing, confocal microscopy, and tank experiments, our data suggest that *L. pedicellatus* and/or *T. bartschi* do not acquire their endosymbiotic bacteria vertically but rather through horizontal transmission. Our findings support a model whereby brooding larvae in adult gill tissue and veliger larvae are aposymbiotic and acquire their symbionts at some point during settlement on wood and/or as they begin burrowing and metamorphosis.

Horizontal acquisition of gill symbionts is consistent with work predicting that *T. turnerae* is a facultative rather than obligate intracellular symbiont. *T. turnerae* lacks many features associated with an obligate intracellular life style such as reduced genome size, skewed GC content, and loss of core metabolic genes [22]. Further, *T. turnerae* has characteristics consistent with a free-living lifestyle including mechanisms to defend against bacteriophage, production of secondary metabolites and the ability to be cultivated *in vitro* without added vitamins or growth factors [20, 22]. The gene content of *T. turnerae* may provide a hint as to its possible environmental niches outside of the host. For example, *T. turnerae* appears to be a specialist for woody plant materials as it has extensive cellulolytic capabilities but lacks enzymes associated with common marine polysaccharides such as agar, alginate, and fucoidan [22]. Given that *L. pedicellatus* and/or *T. bartschi* appear to acquire their bacterial symbionts during association with wooden baits, it is possible that larvae acquire *T. turnerae* from the wood surface either prior to or during initial burrowing. Future work will need to explore whether *T. turnerae* are capable of surviving on the surface of submerged wood in the absence of a shipworm host and/or planktonically in the water column.

Although we were unable to determine the specific environmental source of horizontally acquired shipworm symbionts, our data suggest that juvenile shipworms may acquire them through indirect horizontal transmission. Indirect horizontal transmission occurs when a symbiont is transferred from a host to the environment and then another host [30], and has been observed in other marine symbioses. For example, in the horizontally acquired symbiosis between the *Euprymna scolopes* squid and bioluminescent bacterium *Vibrio fischeri*, adult squid release symbiotically competent bacteria into the water column daily, both seeding the environment with potential symbionts for the next generation and controlling symbiont growth [31–33]. In corals, which primarily acquire their algal symbionts horizontally [34], proliferating *Symbiodinium* cells are preferentially expelled over nonproliferating cells as a way to control symbiotic populations [35] and may also provide a pool of symbionts for coral recruits. Such a phenomenon may also occur for *L. pedicellatus* and/or *T. bartschi*, possibly serving to control symbiont growth within gill tissue and provide juveniles with a pool of potential symbionts.

Our findings contrast previous work describing that a different species of shipworm, *Bankia setacea*, acquires at least one bacterial symbiont through vertical transmission [21]. Shipworms are an incredibly diverse group of marine bivalves with unique communities of gill symbionts [2, 15, 20, 36] and varied reproduction strategies from broadcast spawning to brooding. *B. setacea* is a broadcast spawner, releasing gametes into the water column where they fully develop, which enables a broad distribution of larval settlement [17, 37]. In contrast, *L. pedicellatus* and *T. bartschi* are a long-term brooders, allowing their larvae to fully develop within brood pouches in the gills [38]; *L. pedicellatus* and *T. bartschi* larvae released into the water column rapidly settle within hours and begin boring into wood and metamorphose within days [39–41]. While vertical transmission ensures juvenile *B. setacea* can settle uncolonized wood, potentially reducing competition between parents and progeny [21], horizontal transmission may increase the diversity of symbionts within a shipworm population and favor selection acting on symbionts to benefit their hosts [42]. Notably, the apparent correlation between reproductive strategy and symbiont transmission mode in shipworms is the inverse of observations of Scleractinian corals [5]. In stony corals horizontal transmission is prevalent in broadcast spawners, allowing highly dispersive larvae that can travel long distances to sample from their local environment, while vertical transmission is common in brooders [43, 44]. Additional studies are required to determine whether this trend is apparent across diverse Teredinidae and to elucidate the ecological and evolutionary benefits of horizontal symbiont transmission for *L. pedicellatus and/or T. bartschi*.

Because this work focused on the mode of transmission for *L. pedicellatus and/or T. bartschi* gill symbionts, the composition of such communities across ontogeny remains unknown. Although we were able to culture bacteria consistent with *T. turnerae* physiology [45] from burrowing and more developed juvenile shipworms, it is unclear whether they are the only member of the community at this stage, or whether the community is more complex. Previous work identified at least four distinct bacterial genotypes, including *T. turnerae*, within adult *L. pedicellatus* gill bacteriocytes [15]. Therefore, further investigation should focus on the composition of juvenile *L. pedicellatus and/or T. bartschi* microbiomes throughout morphogenesis and into adulthood, and whether symbiont acquisition is an acute event [46] or continues to occur throughout the shipworm lifespan, as is the case with other marine bivalves [47].

## METHODS

### Cultivation and collection of *L. pedicellatus and/or T. bartschi*

A colony of *L. pedicellatus and T. bartschi* shipworms were collected from found wood in mangroves in the Banana River north of Kelly Park, Merit Island on January 24, 2020 and on the Pineda Causeway on January 25, 2020 and maintained at the Ocean Genome Legacy Center at Northeaster’s Marine Science Center in Nahant, MA. Animals were kept between 25 and 27 C in natural seawater from Nahant at a salinity of 31 ppt.

Adult animals, burrowing juveniles, and more developed juveniles were removed from wooden baits by hand, transferred to filter sterilized autoclaved sea water (FSASW) (0.2 um pore-size filter) and processed (fixed in 4% paraformaldehyde (4% PFA), dissected, or homogenized) within two hours. Veliger larvae were collected via plastic transfer pipettes, transferred to filter sterilized sea water and processed within two hours. Brooding larvae were dissected from gravid adults, rinsed in FSASW, and processed within two hours.

### Symbiont culture quantification

Freshly collected specimens were rinsed with FSASW five times and either processed immediately or frozen at −80°C for at least 24 hours and then processed. Tissue was homogenized using 1.5mL tubes and disposable homogenizer pestles. Tissue homogenate was then serially diluted and plated onto SBM agar plates. Plates were incubated at room temperature and checked daily for colony forming units (CFUs). Brooding and veliger larvae were pooled into groups of three for homogenization and CFU quantification. Treatments where no CFUs were observed are indicated as having the limit of detection, 1 CFU at 10⁰ dilution.

### Shipworm fixation and Fluorescence *in situ* hybridization (FISH) microscopy

Shipworm specimens were fixed, embedded in paraffin, sectioned, and hybridized as described in Betcher *et al*. [18]. Briefly, freshly collected specimens were fixed in 4% PFA in 0.22 μm filtered seawater and incubated at 4°C for 1-3 hours. Each organism was then washed in a series of increasing ethanol solutions (30%, 50%, and 70% x3) for 10 – 15 minutes. Samples were then decalcified in 5% acetic acid in miliQ shaking at 50 rpm overnight at room temperature. Once fully decalcified, samples were again washed in a series of ethanol solutions (30%, 50%, and 70% x3) for 10 - 15 minutes. Larval and juvenile samples were then embedded in autoclaved 4% low melt agarose in PBS to prevent shearing during sectioning. Agar blocks were allowed to harden at room temperature for at least 20 minutes, removed from the mold, and fixed in 4% PFA and stored at 4°C until paraffin embedding.

Paraffin tissue embedding was performed with an automated tissue processor at Beth Israel Deaconess Medical Center, Boston, MA. Paraffin blocks were then mounted onto the microtome, trimmed, sectioned into 5 um thick sections, and mounted onto glass slides and stored at −20°C.

Samples were deparaffinized, rehydrated, and hybridized at the Northeastern University Marine Science center. Samples were deparaffinized by incubating slides in a series of xylene and ethanol solutions using Coplins jars for 3 minutes / solution: xylene x2, 1:1 xylene:100% ethanol, 100% ethanol x2, 95% ethanol, 70% ethanol, and 50% ethanol. Slides were then rehydrated with nanopure water. Oligonucleotide probe hybridizations were performed using Bact338 (seq), a general bacterial-targeted 16S rRNA probe, or NON338 (seq) as a negative control. Probes were mixed in a 1:9 volume solution (5 ng/ul probe final concentration) with hybridization buffer (35% formamide, 0.9 M NaCl, 20 mM Tris-HCl (pH 8.0), 0.01 % SDS). Slides were incubated with each probe for 90 minutes at 46°C, rinsed with washing buffer (35% formamide, 0.7 M NaCl, 20 mM Tris-HCl (pH 8.0), 0.01% SDS) and incubated in preheated washing buffer at 48°C for 25 minutes. Slides were then rinsed with ice cold miliQ, allowed to air dry in the dark, covered in one drop of antifade and a cover slip and allowed to cure in the dark at room temperature for 24 hours. Slides were visualized on a Zeiss LSM 800 inverted confocal laser scanning microscope at the Institute for Chemical Imaging of Living Systems at Northeastern University. Images were visualized and processed using ImageJ software.

## Acknowledgments

We thank Suzanne L. White and the Beth Israel Deaconess Medical Center, Dr. Alex Lovely, and Marvin Altamia for technical assistance, and the Helmuth Lab for equipment. Lauren Speare was funded as a Simons Foundation Awardee of the Life Sciences Research Foundation. Work in the lab of Daniel Distel was supported by the Gordon and Betty Moore Foundation grant no SMS 9339. The authors declare no conflict of interest.

