## Supplemental Figures for "Cultivation and Fluorescent in situ hybridization suggest that some shipworm species acquire endosymbiotic bacteria through indirect horizontal transmission"

### Slide 1
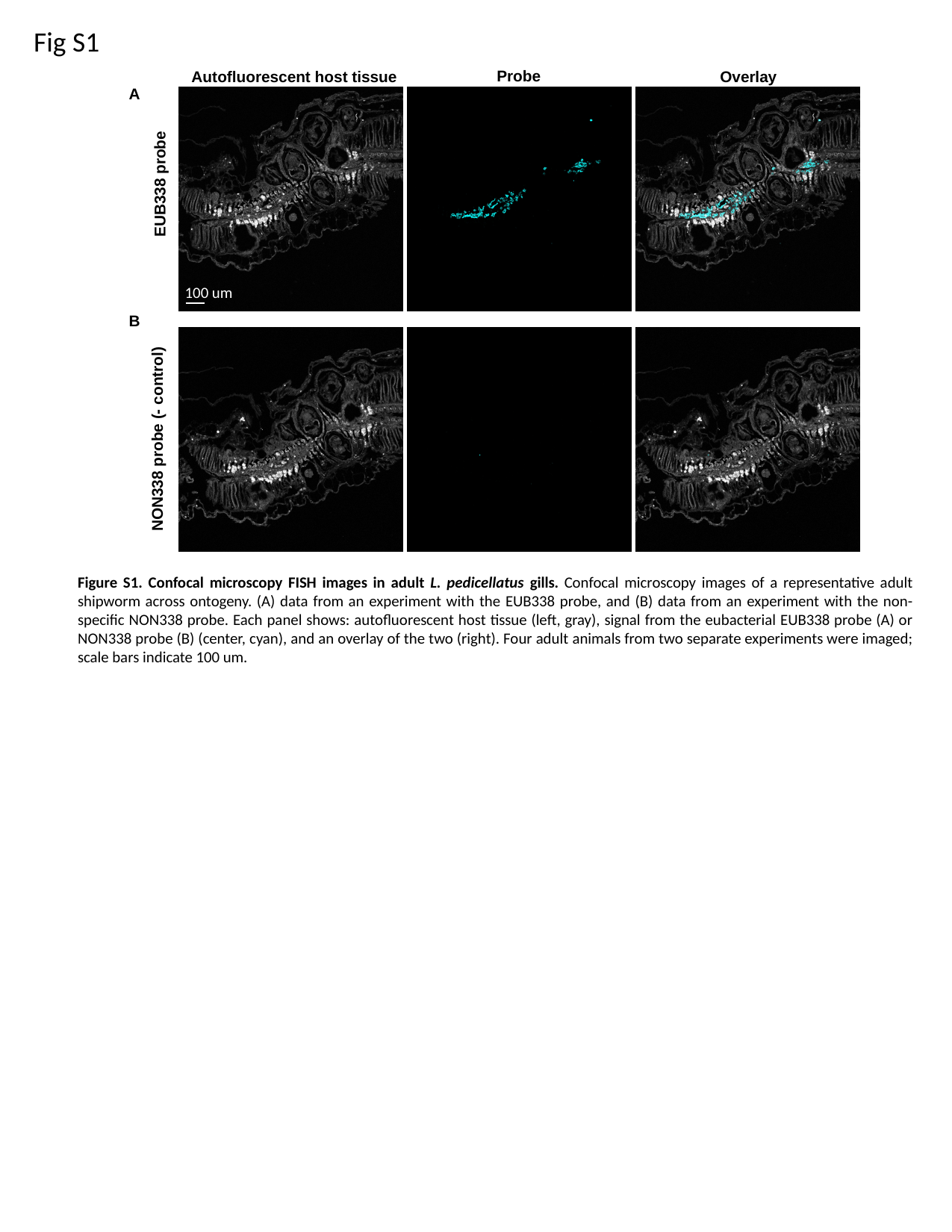

Fig S1
Probe
Overlay
Autofluorescent host tissue
A
B
100 um
EUB338 probe
NON338 probe (- control)
Figure S1. Confocal microscopy FISH images in adult L. pedicellatus gills. Confocal microscopy images of a representative adult shipworm across ontogeny. (A) data from an experiment with the EUB338 probe, and (B) data from an experiment with the non-specific NON338 probe. Each panel shows: autofluorescent host tissue (left, gray), signal from the eubacterial EUB338 probe (A) or NON338 probe (B) (center, cyan), and an overlay of the two (right). Four adult animals from two separate experiments were imaged; scale bars indicate 100 um.

### Slide 2
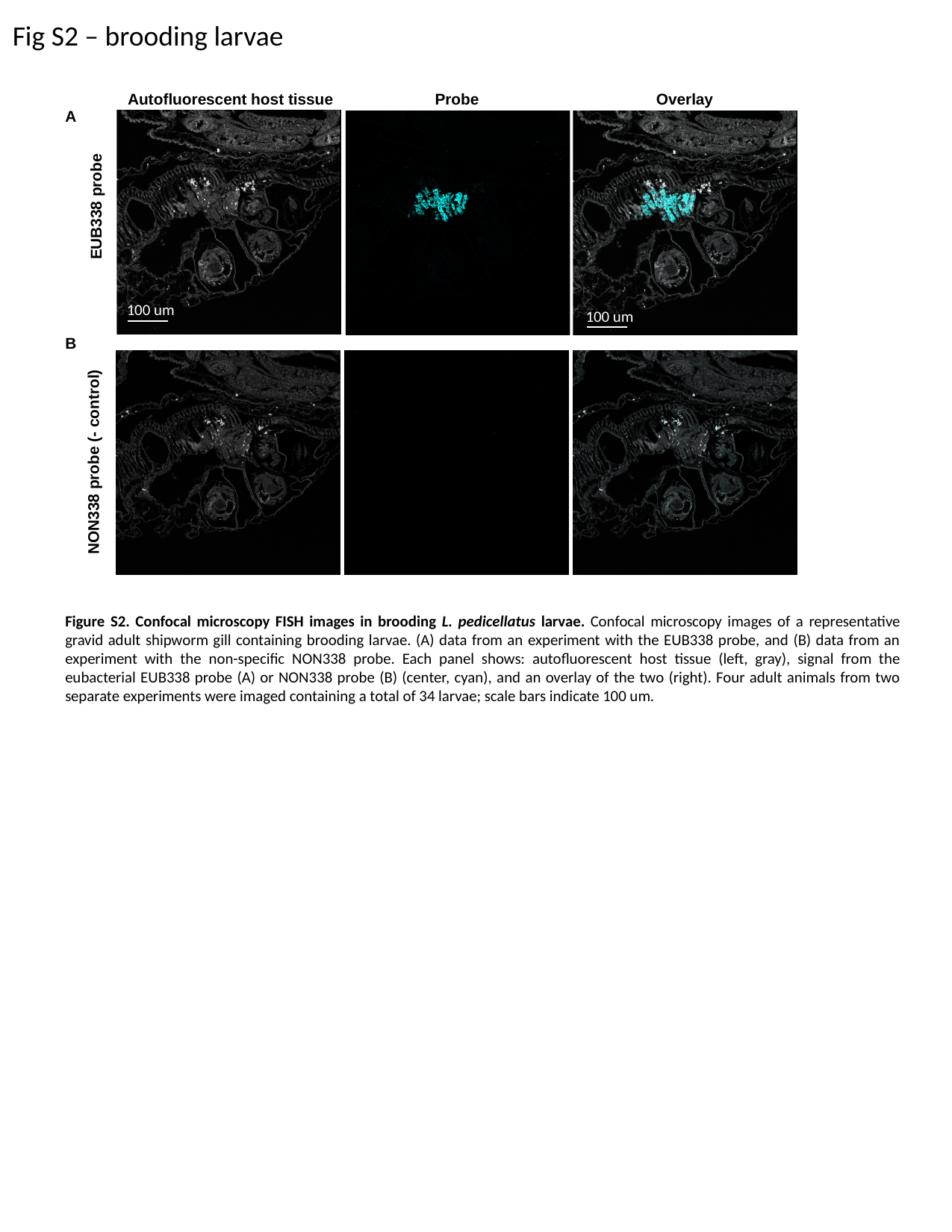

Fig S2 – brooding larvae
Probe
Overlay
Autofluorescent host tissue
A
B
100 um
100 um
EUB338 probe
NON338 probe (- control)
Figure S2. Confocal microscopy FISH images in brooding L. pedicellatus larvae. Confocal microscopy images of a representative gravid adult shipworm gill containing brooding larvae. (A) data from an experiment with the EUB338 probe, and (B) data from an experiment with the non-specific NON338 probe. Each panel shows: autofluorescent host tissue (left, gray), signal from the eubacterial EUB338 probe (A) or NON338 probe (B) (center, cyan), and an overlay of the two (right). Four adult animals from two separate experiments were imaged containing a total of 34 larvae; scale bars indicate 100 um.

### Slide 3
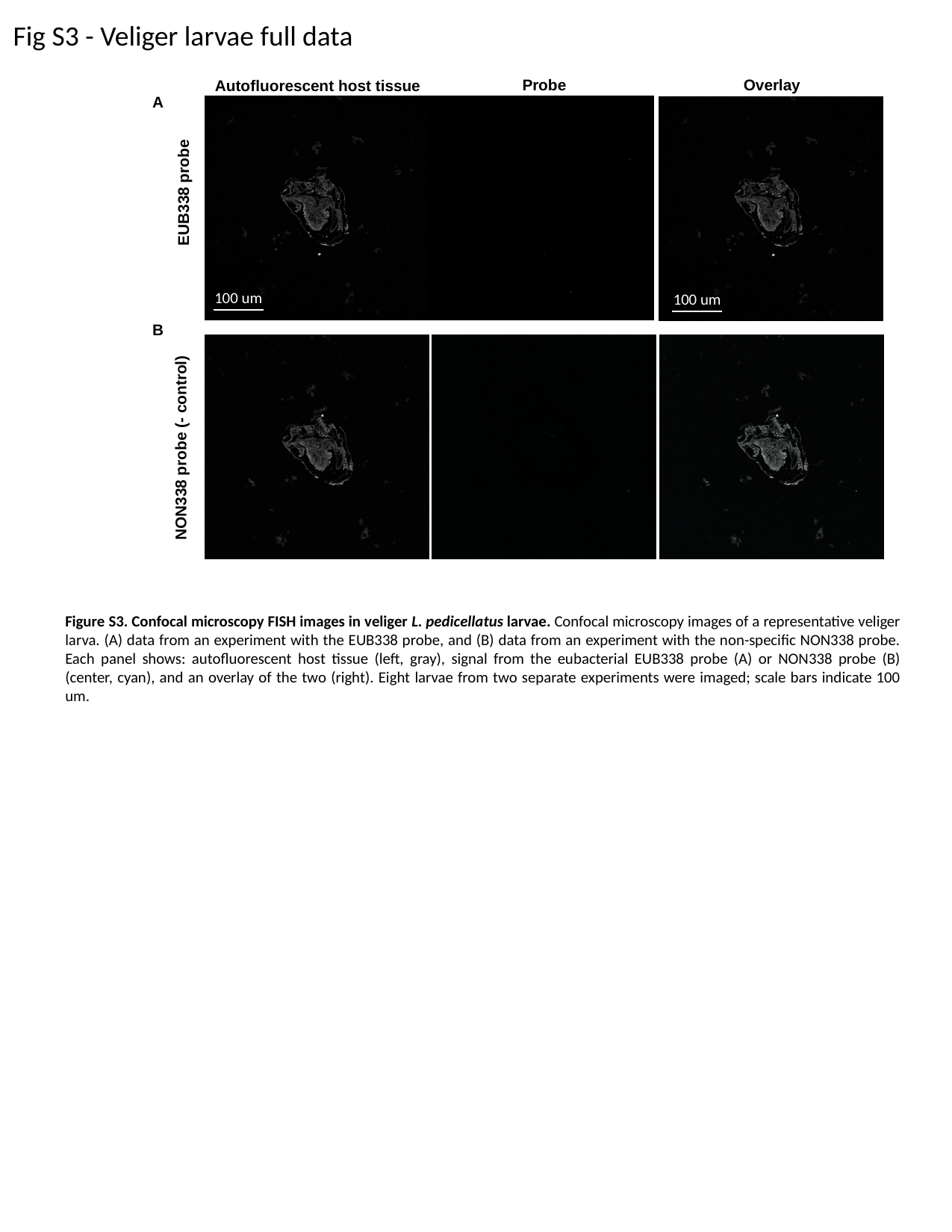

Fig S3 - Veliger larvae full data
Probe
Overlay
Autofluorescent host tissue
A
B
EUB338 probe
NON338 probe (- control)
100 um
100 um
Figure S3. Confocal microscopy FISH images in veliger L. pedicellatus larvae. Confocal microscopy images of a representative veliger larva. (A) data from an experiment with the EUB338 probe, and (B) data from an experiment with the non-specific NON338 probe. Each panel shows: autofluorescent host tissue (left, gray), signal from the eubacterial EUB338 probe (A) or NON338 probe (B) (center, cyan), and an overlay of the two (right). Eight larvae from two separate experiments were imaged; scale bars indicate 100 um.

### Slide 4
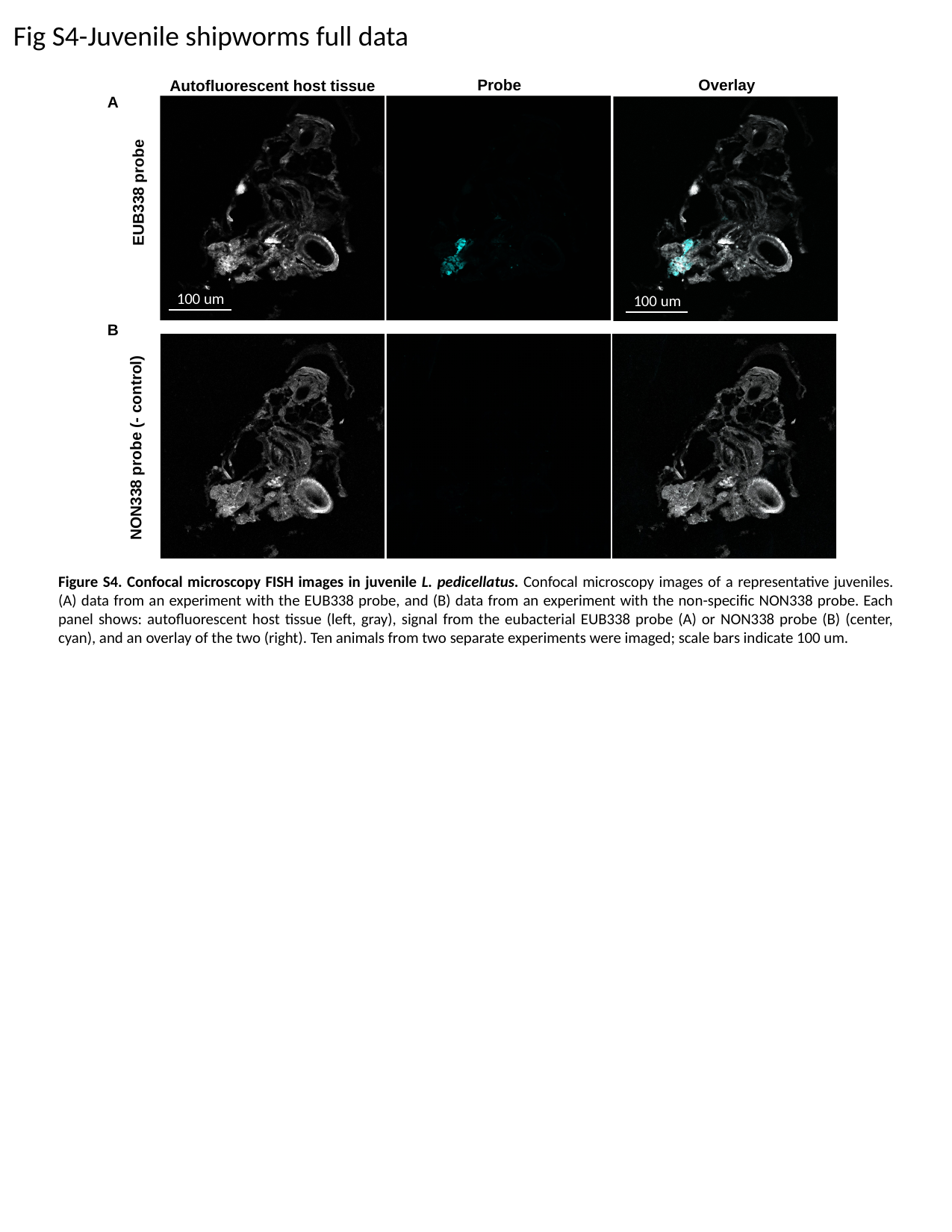

Fig S4-Juvenile shipworms full data
Probe
Overlay
Autofluorescent host tissue
A
B
EUB338 probe
NON338 probe (- control)
100 um
100 um
Figure S4. Confocal microscopy FISH images in juvenile L. pedicellatus. Confocal microscopy images of a representative juveniles. (A) data from an experiment with the EUB338 probe, and (B) data from an experiment with the non-specific NON338 probe. Each panel shows: autofluorescent host tissue (left, gray), signal from the eubacterial EUB338 probe (A) or NON338 probe (B) (center, cyan), and an overlay of the two (right). Ten animals from two separate experiments were imaged; scale bars indicate 100 um.
